# *Muscari commutatum* genome sequences reveal genetic basis of flower pigmentation

**DOI:** 10.64898/2026.09.18.752705

**Authors:** Julie Anne V. S. de Oliveira, Boas Pucker, Tim Böhnert

**Affiliations:** Plant Biotechnology and Bioinformatics, Institute for Cellular and Molecular Botany (IZMB), University of Bonn, Kirschallee 1, 53115 Bonn, Germany; Bonn Institute of Organismic Biology, University of Bonn, Meckenheimer Allee 170, 53115 Bonn, Germany

## Abstract

Flower colour is an important component of floral signaling, yet the genetic basis of naturally occurring colour variation remains poorly understood in many non-model plant groups. Here, we compare genome sequences of a normally pigmented and a largely unpigmented individual of *Muscari commutatum*, a species typically characterized by very dark violet flowers. Both assemblies were approximately 3.5 Gbp in size, with contig N50 values exceeding 130 Mbp and BUSCO completeness of about 99%. Most structural genes of the flavonoid biosynthesis and the corresponding transcriptional regulators were identified in both plants. Strikingly, no flavanone 3-hydroxylase (*F3H*) gene was detected in the genome sequence of the unpigmented plant, and additional manual searches provided no evidence for a bona fide *F3H* copy. Comparison of the two genome sequences revealed a lack of local synteny around the *F3H* locus of the anthocyanin-pigmented plant, suggesting that *F3H* might have been lost in the unpigmented plant in association with a genomic rearrangement.

## Introduction

Floral traits mediate interactions between animal-pollinated plants and their pollinators through combinations of visual, olfactory and structural signals (Raguso, 2004, 2008; Kulahci *et al*., 2008; Schiestl & Johnson, 2013). Flower colour is a particularly conspicuous component of this display and can influence flower detection, recognition and foraging behaviour, although its effects depend on pollinator sensory systems, learning, floral context and background contrast (van der Kooi *et al*., 2019; Trunschke *et al*., 2021). The visual signal produced by a flower is determined not only by pigment identity but also by pigment concentration, tissue structure, and the resulting spectral contrast with the surrounding background (van der Kooi et al., 2019). Accordingly, changes in pigmentation can alter an important component of plant–pollinator signalling, but do not necessarily translate directly into changes in pollinator-mediated selection (Rausher, 2008; Trunschke *et al*., 2021). Identifying the genetic changes underlying naturally occurring colour variation is therefore an important step towards understanding how such floral signals originate and evolve. This is particularly relevant because conspicuous shifts in floral signals can sometimes have a surprisingly simple molecular basis (Schiestl & Johnson, 2013).

Grape hyacinths (*Muscari* Mill.) provide a suitable system for investigating such variation. Their flowers share a relatively conserved basic architecture, with six largely or completely connate tepals forming an urceolate to tubular perianth that is commonly constricted towards the mouth, whereas flower colour varies extensively among species, from yellowish, greenish and brownish tones to pale blue, intense violet, and almost black (Böhnert *et al*., 2023; Hall *et al*., 2025). Despite this diversity, the pollination and reproductive biology of the group has been investigated in comparatively few species (Herrmann *et al*., 2006; Hornemann *et al*., 2012; Canale *et al*., 2014; Çon & Çiçek, 2025). Available studies nevertheless indicate predominantly insect-mediated pollination, involving bees and, to a lesser extent, dipterans, while mating systems can range from mixed mating to predominant or obligate outcrossing (Hornemann *et al*., 2012; Canale *et al*., 2014; Çon & Çiçek, 2025). Experimental evidence nevertheless shows that visual components of the floral display can affect pollinator attraction: the conspicuous sterile flowers of *Leopoldia comosa* (= *Muscari comosum*) strongly increased pollinator attraction and reproductive performance, whereas the much less conspicuous sterile flowers of *M. armeniacum* had no detectable effect on visitation or reproductive success (Morales *et al*., 2013; Gavini, 2025). These studies demonstrate that visually conspicuous floral traits can influence pollinator interactions in *Muscari*, providing a functional context for investigating the pronounced flower-colour variation within the genus, even though the role of colour itself remains largely unexplored.

*Muscari commutatum* Guss., a species of the central and eastern Mediterranean extending from Italy through the Balkan Peninsula and Greece to southwestern Türkiye (Uysal *et al*., 2021; Böhnert *et al*., 2023), lies near one extreme of this colour spectrum: its fertile flowers are typically dark violet to blackish-violet (Vladimirov, 2022; Böhnert *et al*., 2023). Rare plants, however, show an almost complete loss of this pigmentation, which have been observed independently at different localities within the species’ range (Vladimirov, 2022). On Lesvos, ivory-white to pale green or pinkish individuals have been recorded at several sites, including one population with approximately 50 depigmented plants growing together with about 200 normally pigmented individuals (Vladimirov, 2022). The unpigmented individual investigated here originated from the Peloponnese (Greece), providing an independent occurrence of the same conspicuous phenotype. Comparable colour-loss phenotypes in other *Muscari* species have been linked to changes in anthocyanin metabolism and its transcriptional regulation (Lou *et al*., 2014; Ma *et al*., 2023). The extreme phenotypic contrast between these forms makes *M. commutatum* particularly suitable for identifying genomic changes associated with floral depigmentation and provides a molecular basis for subsequently asking how such variation arises, persists and potentially affects floral signalling. At the biochemical level, blue and violet flower coloration in *Muscari* is largely associated with anthocyanins, particularly delphinidin-derived pigments (Yoshida *et al*., 2009; Lou *et al*., 2014, 2017), which form part of the broader flavonoid pathway.

Flavonoids are a large group of specialized metabolites derived from the aromatic amino acid phenylalanine (Winkel-Shirley, 2001). Flavonols, flavones, anthocyanins, and proanthocyanidins are the major compound groups that are produced by different branches of the flavonoid biosynthesis (Winkel-Shirley, 2001). Flavonoids fulfill a variety of physiological and ecological functions in plants, many of which are connected to their antioxidant properties (Grotewold, 2006). For example, flavonols are well known for their involvement in UV response (Stracke *et al*., 2007; Owens *et al*., 2008a; Stracke *et al*., 2010). Proanthocyanidins are involved in seed coat pigmentation, which has been extensively studied in *Arabidopsis thaliana* (Sagasser *et al*., 2002; Buer & Djordjevic, 2009; Appelhagen *et al*., 2014). Among the products of flavonoid biosynthesis, anthocyanins are the most noticeable group due to their bright colors. Many flower and fruit colors are caused by the accumulation of anthocyanins, which can confer a variety of hues including red, pink, purple, and blue (Grünig *et al*., 2025). The specific color is determined by the chemical structure of the anthocyanin, with hydroxylation at the B-ring playing a fundamental role: one hydroxy group results in orange, two hydroxy groups result in magenta, and three hydroxy groups lead to blue (Seitz *et al*., 2006; Schwinn *et al*., 2014). Additional decoration of anthocyanins with sugars, acyl groups, or methyl groups further influences the molecule’s properties, including the color (Grünig *et al*., 2025). Anthocyanins are considered important for the attraction of pollinators and seed dispersers (Davies *et al*., 2012; Grünig *et al*., 2025), but also play important roles in response to biotic and abiotic stresses (Jezek *et al*., 2023; Muralidhar *et al*., 2026). Despite 200 years of research on anthocyanins, the relative importance of the different functions is still an active field of research and might be lineage-specific (Choudhary *et al*., 2026b). The presence of anthocyanins in most land plants has resulted in the assumption that this pigment biosynthesis pathway would be highly conserved, but this notion has recently been challenged by the discovery of lineage-specific differences (Khatun *et al*., 2025; Choudhary *et al*., 2026a). Fundamental discoveries might still be possible in a biosynthesis pathway that has been widely considered a model system for specialized metabolism in plants.

The biosynthesis of anthocyanins branches from the general flavonoid pathway. Chalcone synthase (CHS), chalcone isomerase (CHI), and flavanone 3-hydroxylase (F3H) are required for producing dihydroflavonols, which serve as precursors for the anthocyanin biosynthesis (Winkel-Shirley, 2001; Grotewold, 2006). The specific enzymes involved in anthocyanins are under debate because large parts of the anthocyanin biosynthesis overlap with the proanthocyanidin biosynthesis (Choudhary *et al*., 2026b). Dihydroflavonol 4-reductase (DFR), anthocyanidin synthase (ANS), anthocyanin-related glutathione S-transferase (arGST), and UDP-dependent anthocyanidin 3-O-glycosyltransferase (UGT) are generally considered as anthocyanin biosynthesis enzymes (Pelletier *et al*., 1997; Saito *et al*., 1999; Johnson *et al*., 2001; Tohge *et al*., 2005; Eichenberger *et al*., 2023; Choudhary *et al*., 2026b). Decoration enzymes can add additional sugar moieties, acyl groups, or methyl groups, resulting in a plethora of chemically different anthocyanin derivatives (Tohge *et al*., 2005; Luo *et al*., 2007; Yonekura-Sakakibara *et al*., 2012; Kovinich *et al*., 2014; Grünig *et al*., 2025).

The biosynthesis of anthocyanins takes place at the endoplasmic reticulum (ER), and there is evidence for a metabolon formed by the involved enzymes (Winkel-Shirley, 1999; Nakayama *et al*., 2019). Since anthocyanins are stored in the central vacuole, transport from the site of biosynthesis to the storage location is required. There are currently two models that could explain these observations: (1) transport through the cytoplasm and import into the vacuole over the tonoplast or (2) direct import into the ER lumen and transport in vesicles towards the central vacuole (Goodman *et al*., 2004; Poustka *et al*., 2007; Kaur *et al*., 2021; Pucker & Selmar, 2022; Grünig *et al*., 2025).

The anthocyanin biosynthesis is controlled at the transcriptional level by several transcription factors that act in concert (Grünig *et al*., 2025). Biosynthesis genes are activated by the MBW complex, comprising a MYB protein, a bHLH protein, and a WD40 protein (Ramsay & Glover, 2005; Gonzalez *et al*., 2008; Li, 2014; Lloyd *et al*., 2017). There are also negative regulators that can interfere with the MBW complex or target the anthocyanin biosynthesis genes directly (LaFountain & Yuan, 2021). While anthocyanins have been reported in response to multiple stresses, it appears that the anthocyanin biosynthesis in many plants is particularly induced by light (Ma *et al*., 2021; Nowak *et al*., 2024; Meckoni *et al*., 2026).

Anthocyanins have been investigated for about 200 years, facilitated by the visible phenotypes that make the identification and investigation of mutants easy (McClintock, 1950; Marin-Recinos & Pucker, 2024; Choudhary *et al*., 2026a). While the linear part of the anthocyanin biosynthesis could be blocked at any step by disrupting mutations in structural genes, previous studies indicated that the block is often caused by changes in the anthocyanin biosynthesis-activating transcription factors, especially the anthocyanin-specific MYBs (Marin-Recinos & Pucker, 2024). Due to the pleiotropic role of many involved genes, changes in this anthocyanin-specific MYB might be associated with a weaker evolutionary disadvantage. While anthocyanin loss has been frequently reported as intraspecific variation (Marin-Recinos & Pucker, 2024), reports of anthocyanin loss at the family level are limited to some Caryophyllales and Cucurbitaceae (Mabry, 1964; Stafford, 1994; Pucker *et al*., 2024; Choudhary *et al*., 2026a).

Here, we set out to unravel the genetic basis of the intraspecific pigmentation variation in *M. commutatum* in order to support taxonomic studies. Given that differences in flower pigmentation can support speciation events, we ask whether the investigated plants are still members of the same species.

## Materials and Methods

### DNA extraction and nanopore sequencing

*Muscari commutatum* samples were collected from the Botanical Gardens of the University of Bonn, a representative of a pigmented plant: GR-0-BONN-33000, and an unpigmented plant: GR-0-BONN-47263. The leaves were harvested and homogenized by grinding in liquid nitrogen; high-molecular-weight DNA was extracted with a modified CTAB-based protocol as previously described (Siadjeu *et al*., 2020). Next, quality control of the DNA was performed, initially via NanoDrop measurement, followed by agarose gel electrophoresis, and Qubit measurement as previously described (de Oliveira *et al*., 2026). Short DNA fragments were then depleted with the Short Read Eliminator kit (Pacific Biosciences) following the supplier’s instructions. Libraries for nanopore sequencing were prepared with the SQK-LSK114 ligation-based kit (Oxford Nanopore Technologies, ONT) using 1 µg of DNA and following the supplier’s instructions. Sequencing was performed on a PromethION 2 Solo with R10.4.1 flow cells. Due to blockage of a large proportion of pores, a wash step was performed before the loading of a new library to achieve optimal performance of the flow cells. Basecalling of the raw sequencing data was performed with Dorado v1.3.1 (ONT) with a high-accuracy model (dna_r10.4.1_e8.2_400bps_hac@v5.2.0) on an NVIDIA L4 GPU in the de.NBI cloud.

### Genome sequence assembly and annotation

Both *M. commutatum* genome sequences were assembled with Hifiasm-0.25.0-r726 (Cheng *et al*., 2021). Assembly statistics were calculated via contig_stats3.py (de Oliveira *et al*., 2026), and contig names were cleaned to avoid technical issues in the downstream analysis using clean_genomic_fasta.py (de Oliveira *et al*., 2026). Assembly completeness was assessed using BUSCO v6.0.0 (Tegenfeldt *et al*., 2025) in the genome mode based on the liliopsida_odb12 reference dataset. For the structural annotation, RNA-seq datasets (Additional file 1) were retrieved from the Sequence Read Archive using ‘fastq-dump’. The RNA-seq read mapping was generated with HISAT2 v2.2.1, and the resulting BAM file was used for GeMoMa v1.9 (Keilwagen *et al*., 2019), which enables the inference of hints for gene prediction. The following datasets from were used as hints: *Asparagus officinalis* (GCF_001876935.1) (Harkess *et al*., 2017), *Iris pallida* (GCA_029216955.1) (Bruccoleri *et al*., 2023), *Platanthera zijinensis* (GCA_039513925.1) (Li *et al*., 2022), *Dendrobium nobile* (GCA_022539455.1) (Xu *et al*., 2022), *Dendrobium chrysotoxum* (GCA_019925795.1) (Zhang *et al*., 2021), *Dendrobium thyrsiflorum* (GCA_040670175.1) (Chen *et al*., 2024), *Platanthera guangdongensis* (GCA_039583875.1) (Li *et al*., 2022), *Apostasia shenzhenica* (GCA_002786265.1) (Zhang *et al*., 2017), *Dendrobium catenatum* (GCF_001605985.2) (Zhang *et al*., 2016), and *Phalaenopsis equestris* (GCF_001263595.1) (Cai *et al*., 2015).

The initial annotation was filtered with GeMoMa using the following criteria: f=“start==‘M’ and stop==‘*’ and (score/aa>=‘0.75’)”; the RNA-seq evidence was prepared with GeMoMa’s ‘ERE’ based on the read mapping generated with HISAT2, followed by an analysis of all predicted polypeptide sequences with BUSCO v6.0.0 with the liliopsida_odb12 lineage, in the protein mode to assess the annotation completeness. Another filtering step was done with: f=“start==‘M’ and stop==‘*’ and aa>=15 and (isNaN(score) or score/aa>=‘1.5’)” atf=“tie==1 or sumWeight>1”, and the resulting predicted genes were renamed with ‘AnnotationFinalizer’, and CDS and polypeptide sequences were extracted with GeMoMa’s ‘Extractor’. Functional annotation terms were assigned to the predicted polypeptide sequences with the Python script construct_anno.py (Pucker & Iorizzo, 2023) based on the Araport11 annotation of *Arabidopsis thaliana* (Cheng *et al*., 2017).

### Investigation of the flavonoid biosynthesis

Genes involved in flavonoid biosynthesis were annotated by KIPEs v3.2.7 (Rempel *et al*., 2023) using the flavonoid bait set v3.4. The anthocyanin biosynthesis-activating MYB transcription factors were identified with the MYB_annotator v1.0.3 running with default parameters (Pucker, 2022). The anthocyanin biosynthesis-activating bHLH protein TT8 was identified with the bHLH_annotator v1.04 running with default parameters (Thoben & Pucker, 2023). TTG1 was identified based on a collection of orthologous sequences using collect_best_BLAST_hits.py v0.36 (Pucker & Iorizzo, 2023) to find candidate sequences in *M. commutatum*. The TTG1 bait sequences and the newly identified candidate sequences were subjected to MAFFT v7 (Katoh & Standley, 2013) for construction of a global alignment. IQ-TREE3 v3.1.8 (Wong *et al*., 2026) was used to infer a gene tree from the alignment with the parameters -alrt 1000 -bb 1000 and the Q.PLANT+F+I+R4 model. Manual tree inspection was conducted via iTOL v6 (Letunic & Bork, 2024).

A manual search for F3H candidates in the genome sequence of the unpigmented plant was conducted with tBLASTn v2.13.0+ (Altschul *et al*., 1990; Camacho *et al*., 2009) using an e-value cutoff of 0.000001.

## Results & Discussion

### *Muscari commutatum* genome sequences and annotation

The genome sequences of the pigmented and unpigmented *Muscari commutatum* were assembled based on ONT long-read sequencing data with Hifiasm (**Table 1**). The assembly sizes of 3.5 Gbp for both plants are consistent with results of a flow cytometry study in *Muscari*, which reported a mean DNA amount of 4.18 pg for 1C (https://cvalues.science.kew.org). The pigmented assembly comprised 265 contigs, with an N50 of 132 Mbp and a GC content of 44%, indicating a high level of contiguity. Assembly completeness was estimated at 99% based on BUSCO genes (liliopsida_odb12). For the unpigmented plant, the assembly comprised 238 contigs, with 148 Mbp N50 and 44% GC content; assembly completeness was estimated at 98.9% based on BUSCO using the same lineage.

**Table 1.** Statistics of representative genome sequences generated based on *M. commutatum* nanopore sequencing data of an anthocyanin-pigmented and an unpigmented plant, respectively.

| Hifiasm | Anthocyanin-pigmented plant | Unpigmented plant |
| --- | --- | --- |
| Assembly size | 3.5 Gbp | 3.5 Gbp |
| Number of contigs | 265 | 238 |
| N50 | 132 Mbp | 148 Mbp |
| BUSCO | C:99.0%[S:83.9%,D:15.1%],<br>F:0.4%,M:0.6%,n:2821 | C:98.9%[S:82.9%,D:16.0%],<br>F:0.5%,M:0.6%,n:2821 |

The structural annotation of all protein-encoding genes helps with studies of the genetic basis of flower color formation. GeMoMa generated the best structural annotation using hints from several plant species (**Table 2**). A completeness check of the annotation with BUSCO revealed about 82% of all expected genes in both plants. Unfortunately, there are no references of closely related species to *Muscari commutatum* for the structural annotation, therefore, not all expected BUSCO genes were discovered.

**Table 2.** Structural annotation of anthocyanin-pigmented and unpigmented *Muscari commutatum*. GeMoMa was supplied with RNA-seq hints and data sets of *Asparagus officinalis* (GCF_001876935.1), *Iris pallida* (GCA_029216955.1), *Platanthera zijinensis* (GCA_039513925.1), *Dendrobium nobile* (GCA_022539455.1), *Dendrobium chrysotoxum* (GCA_019925795.1), *Dendrobium thyrsiflorum* (GCA_040670175.1), Platanthera *guangdongensis* (GCA_039583875.1), *Apostasia shenzhenica* (GCA_002786265.1), *Dendrobium catenatum* (GCF_001605985.2), and *Phalaenopsis equestris* (GCF_001263595.1).

|  | Anthocyanin-pigmented | Unpigmented |
| --- | --- | --- |
| **Number of genes** | 51066 | 50783 |
| **Number of transcripts** | 79760 | 79753 |
| **BUSCO** | C:81.7%[S:48.8%,D:32.9%],<br>F:11.8%,M:6.5%,n:2821 | C:81.5%[S:48.5%,D:33.0%],<br>F:11.7%,M:6.8%,n:2821 |

### Anthocyanin biosynthesis genes in Muscari

All genes required for the biosynthesis of anthocyanins have been identified in the genome sequence of the anthocyanin-pigmented plant (**Fig. 2**). This includes structural genes encoding CHS, CHI, F3H, F3’H, DFR, ANS, arGST, and UGT, as well as genes encoding the corresponding transcriptional regulators arMYB, the bHLH protein TT8, and the WD40 protein TTG1. FLS, required for the flavonol biosynthesis, which competes with the anthocyanin biosynthesis for dihydroflavonols as substrates, was detected. No LAR was identified, which aligns with observations in a number of plant lineages that also lack LAR (Wang *et al*., 2018; Marin-Recinos & Pucker, 2025). The results regarding ANR are inconclusive because some amino acid residues that are generally considered important for the functionality are absent. However, it is plausible that lineage-specific differences in *Muscari* are responsible for this observation. There is currently no evidence for differences in the flavonol or proanthocyanidin biosynthesis branch between the anthocyanin-pigmented and the unpigmented plant investigated in this study.

**Fig. 1.**
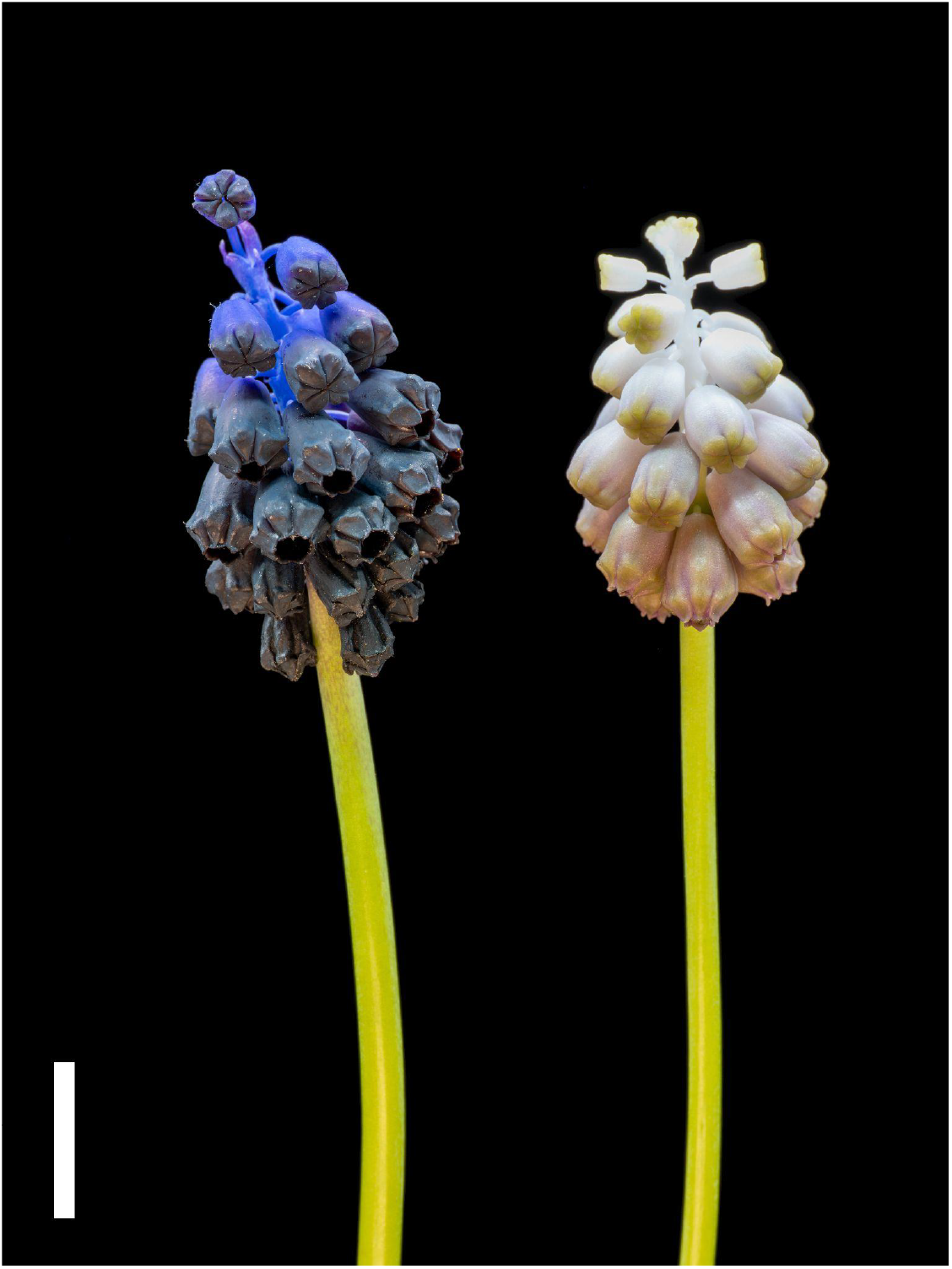
Pictures of *Muscari commutatum*. Anthocyanin-pigmented plant (left) and largely unpigmented plant (right). The scale bar is 1cm.

**Fig. 2.**
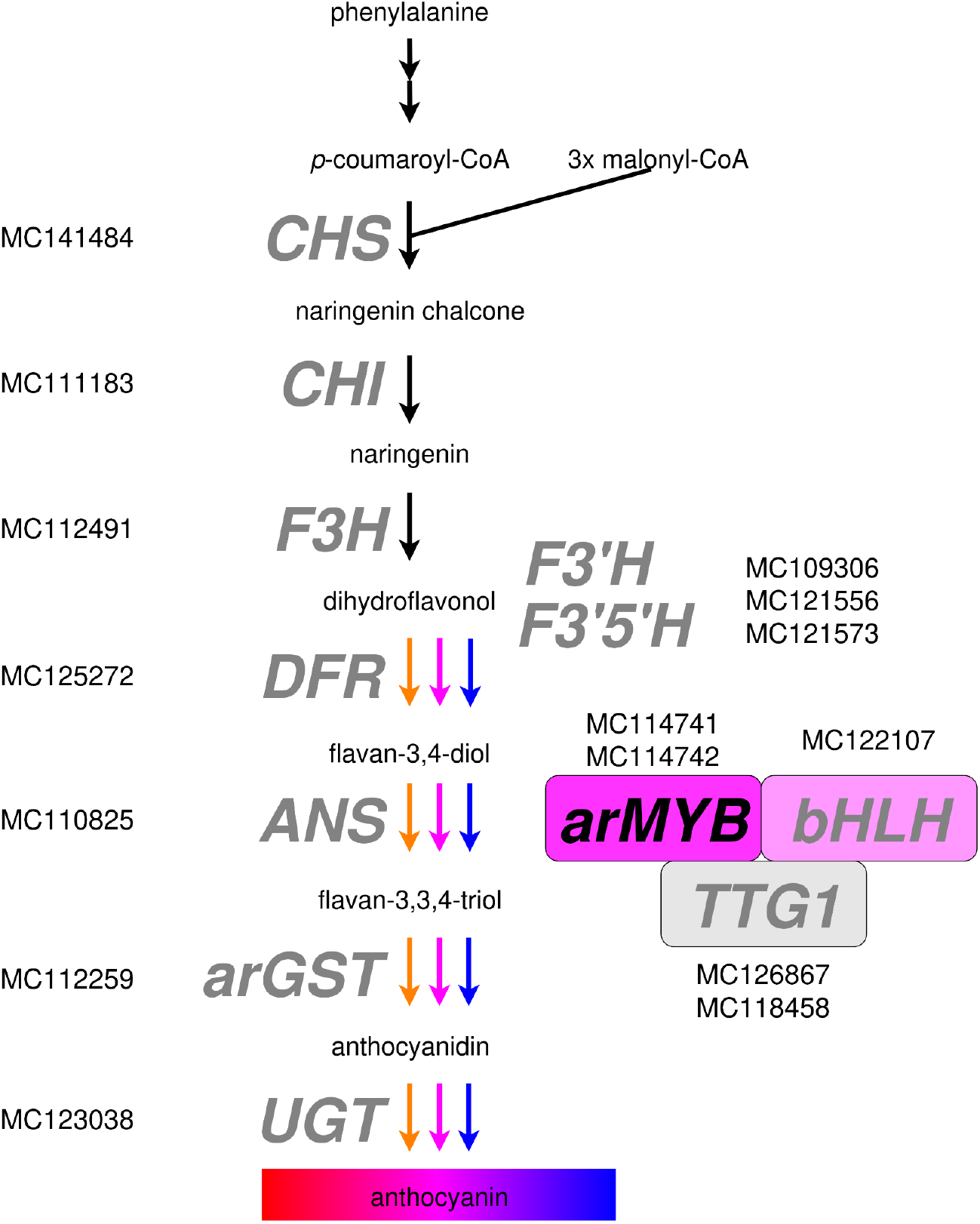
Anthocyanin biosynthesis genes identified in the genome sequence of the anthocyanin-pigmented *M. commutatum* plant. The figure layout was derived from (Horz *et al*., 2026).

While most anthocyanin biosynthesis genes have been discovered in the genome sequence of the unpigmented plant, no *F3H* was annotated. This gene plays a crucial role in the biosynthesis of flavonols, anthocyanins, and proanthocyanidins (Pelletier & Shirley, 1996; Owens *et al*., 2008b). This pleiotropic role should result in a strong selection against the loss of *F3H*, but mutants have been reported (Pelletier & Shirley, 1996). To rule out an annotation artifact as an explanation for the absence of *F3H*, a manual search with the *M. commutatum* F3H sequence against the genome sequence of the unpigmented plant was conducted, but it only revealed hits with about 85% sequence similarity. Since *F3H* belongs to the large gene family of 2-oxoglutarate dioxygenases, this is most likely another member of the family, but not a *bona fide F3H*. This suggests that *F3H* is not represented in this genome sequence. Given the high completeness of the genome sequence (99% of BUSCO genes detected), it appears very unlikely that the absence of *F3H* would be the result of an assembly artifact.

A frequently applied approach to support hypotheses about the absence of a gene is a synteny analysis around the locus where a gene or a group of genes would be expected to demonstrate that the locus is “empty” (Pucker *et al*., 2024; Khatun *et al*., 2025; Choudhary *et al*., 2026a). A synteny analysis was attempted based on the *F3H* locus in the genome sequence of the anthocyanin-pigmented plant. However, genes flanking *F3H* appeared not to be syntenic between the two *M. commutatum* plants investigated in this study. This suggests large-scale genomic rearrangements as an explanation for the observed differences in gene order. It has not escaped our notice that such rearrangements could be associated with the absence of *F3H* in the unpigmented plant.

Like FLS, FNSII, and ANS, F3H is a member of the 2-oxoglutarate dioxygenase family that has been attributed with bifunctionality (Park *et al*., 2019; Schilbert *et al*., 2021). Therefore, it is not possible to rule out the presence of F3H activity in *M. commutatum*, because FLS or ANS might harbour some F3H side activity. Future studies of the different 2-ODD members in *Muscari* are required to unravel the specific enzymatic capabilities of each family member.

Taken together, our results identify loss of *F3H*, potentially associated with structural genomic rearrangement, as a plausible molecular basis for the strongly depigmented phenotype observed in *M. commutatum*. The broader evolutionary significance of such colour-loss variants remains unresolved. Depigmented individuals have been documented in geographically separated parts of the species’ range, including the Peloponnese and Lesvos, where they may occur locally at appreciable frequencies (Vladimirov, 2022). Similar pale- or white-flowered phenotypes also occur in other *Muscari* lineages, including *M. vanensis*, which is distinguished as a separate species by a combination of morphological, karyological, molecular and palynological characters rather than flower colour alone (Uysal *et al*., 2022). These observations illustrate that conspicuous changes in floral pigmentation can occur at different levels of biological differentiation, while their molecular origin and evolutionary persistence may be independent questions. Population-level sampling will therefore be required to determine whether depigmentation in *M. commutatum* has arisen repeatedly, whether similar alleles are shared among populations, and whether these colour variants have consequences for gene flow, reproductive success or pollinator interactions.

## Declarations

### Ethics approval and consent to participate

Not applicable

### Consent for publication

Not applicable

### Availability of data and materials

All datasets associated with this study are publicly available. The genome sequence and corresponding annotation are available via bonndata (https://doi.org/10.60507/FK2/G2TRGZ).

### Authors’ contributions

TB and BP conceived the study. TB provided the plant material. JAVSdO conducted the DNA extraction, sequencing, and bioinformatic analyses. BP supervised the work, conducted bioinformatic analyses, and wrote the manuscript, with additions from TB. All authors approved the final version of the manuscript and agreed to its submission.

## Acknowledgements

This work was supported by the de.NBI Cloud within the German Network for Bioinformatics Infrastructure (de.NBI) and ELIXIR-DE (Forschungszentrum Jülich and W-de.NBI-001, W-de.NBI-004, W-de.NBI-008, W-de.NBI-010, W-de.NBI-013, W-de.NBI-014, W-de.NBI-016,

W-de.NBI-022). We thank all members of the Plant Biotechnology and Bioinformatics group for their support and feedback during the process. We are grateful for the excellent support provided by the team of the University of Bonn Botanic Gardens.

